# Taylor’s power law rules the dynamics of allele frequencies during viral evolution in response to host changes

**DOI:** 10.1101/2024.12.27.630372

**Authors:** João M. F. Silva, María J. Olmo-Uceda, Valerie J. Morley, Paul E. Turner, Santiago F. Elena

**Affiliations:** Instituto de Biología Integrativa de Sistemas (I2SysBio), CSIC – Universitat de València, Paterna, Valencia, Spain; Department of Ecology and Evolutionary Biology, Yale University, New Haven, CT, USA; QBio Institute, Yale University, New Haven, CT, USA; The Santa Fe Institute, Santa Fe, NM, USA

**Keywords:** Approximate Wright-Fisher diffusion model, effective population size, experimental evolution, selection coefficient, Sindbis virus, Taylor’s temporal fluctuation scaling, Hurst’s exponent, virus evolution

## Abstract

Sudden and gradual changes in host species composition —transitioning from more permissive to less permissive hosts— have been shown to influence viral fitness, virulence, and rates of molecular evolution. In this study, we analyzed the roles of stochasticity and selection using previously published time-series data from evolving populations of Sindbis virus under different rates of host replacement. First, an approximate Markov model within the Wright-Fisher diffusion framework revealed a reduction in effective population size by approximately half in scenarios involving sudden host changes. These scenarios were also associated with fewer and generally weaker beneficial mutations. Second, genetic distance analysis between populations at consecutive time points indicated that populations undergoing gradual host changes evolved steadily until the original host was no longer present. Distances to the ancestral sequence in these cases exhibited occasional leapfrog phenomena, where the rise of certain haplotypes is not predictable based solely on their relatedness to previously dominant ones. In contrast, populations exposed to sudden host changes displayed less stable compositions and diverged from the ancestral sequence at a more constant rate. Third, we observed that allele frequencies distribution followed a power law consistent with Taylor’s law. Both treatments exhibited high levels of allele aggregation with significant fluctuations, where neutral, beneficial, and deleterious alleles could be distinguished by their general behavior and their position on Taylor’s plot. Finally, we found evidence that the host replacement regime influences the temporal distribution of mutations across the genome.

## 1. Introduction

The molecular dynamics of adaptation are heavily influenced by environmental heterogeneity. Pathogens adapt to new hosts by fixing mutations that enhance fitness in the novel host, even if these mutations are neutral or detrimental in the original host. Consequently, the availability of novel hosts in the environment affects both the rate and direction of molecular adaptation. For example, the adaptation dynamics of the Sindbis virus (SINV) to a less permissive cell type in laboratory tissue culture are determined by the rate at which these novel host cells “invade” the environment [1,2]. In a series of evolution experiments, Morley et al. [1] introduced a novel host cell type at varying rates, ranging from a gradual increase in the proportion of the novel cell type with each passage to an abrupt shift to an environment composed entirely of the novel host. Gradual changes were associated with virus populations achieving higher fitness in both novel and original hosts, as well as greater convergence among populations that fixed the same adaptive mutations [1].

Allele frequencies typically fluctuate over time in both natural and experimental populations due to a variety of factors, including purely stochastic processes (*e.g.*, genetic drift, fluctuating selection, migration, or unpredictable ecological changes) and deterministic processes (*e.g*., directional selection). Indeed, temporal fluctuations are ubiquitous in physical systems, raising the question of whether they follow universal laws. Various power-law relationships have been found to be pervasive in physical and biological systems. One such relationship, known as Taylor’s power law —or the fluctuation scaling law— posits that the variance of a system’s, *σ*^2^, elements scales as a power of the mean, *μ*: *σ*^2^= *Vμ^β^* [3]. Originally described in ecology, this law naturally arises in many complex systems [4–9]. Its parameters capture both the amplitude of the noise level (*V*) and the degree of temporal aggregation (*β*) in the fluctuations observed within the system [5,6,8]. Notably, in the context of infectious diseases, Taylor’s law has been applied to temporal data from the human microbiota, revealing that an individual’s negative health status is associated with increased noise and system instability [8]. Similarly, recent studies have found that during severe acute respiratory syndrome coronavirus 2 (SARS-CoV-2) infection, the transcripts dynamics in cells from human intestinal organoids, but not pulmonary cells, exhibit an increase-decrease-increase pattern in system noise and instability as infection progresses and the virus accumulates [9].

In the context of temporal variations in allele frequencies during virus adaptation to novel hosts, we propose that Taylor’s power law can be used to model how the variance of allele frequencies changes over time. In populations influenced solely by stochastic processes (*i.e*., Exponential), it is expected that *β* = 2, meaning that variance scales quadratically with the mean allele frequency. In contrast, a *β* > 2 indicates nonrandom processes that amplify variability, such as migration or environmental heterogeneity, whereas a *β* < 2 suggests processes that constrain variability, such as stabilizing selection or density dependence (with *β* = 1 corresponding to a Poisson process).

Many natural processes exhibit long-term memory or persistent behavior (autocorrelation), meaning that a high value is likely to be followed by another high value (and similarly for low values) [10–15]. This behavior can be quantified using the Hurst exponent (denoted as *H* throughout this work), where: 0 < *H* < 0.5 indicates anti-persistent (negative autocorrelation) behavior, *H* = 0.5 indicates a random walk, and 0.5 < *H* < 1 indicates persistent behavior. Interestingly, many studied processes have an estimated *H* ≈ 0.7, a phenomenon known as the Hurst phenomenon [10]. Although Taylor’s law and the Hurst phenomenon both describe variability and scaling behavior, they focus on different aspects of the process and are indirectly related. Specifically, when Taylor’s law is applied to the variance of fluctuations at different time scales, the simple relationship *β* = 2*H* must hold [11]. This connection arises because both laws describe the fractal or scaling properties of purely stochastic processes.

Here, we estimate population genetic parameters —such as the selection coefficient per allele (*s*) and effective population size (*N_e_*)— and describe allele dynamics in experimentally evolving SINV populations under two different temporal schemes of host replacement. SINV populations that experienced a sudden replacement of a highly susceptible host with a less permissive one exhibited smaller *N_e_* values, indicating stronger bottlenecks and genetic drift. This observation prompted us to investigate how noise and selection affect the dynamics of virus molecular adaptation to novel hosts. First, we show that under gradual replacement of the highly susceptible host, the genetic composition of the viral populations changed steadily until the host was completely removed. In contrast, under the sudden treatment, the shift in population composition was more pronounced and less consistent. Next, we characterized Taylor’s power law for the temporal variation in allele frequency. This power model allowed us to examine in greater detail how selection and noise resulting from genetic drift influence the dynamics of molecular adaptation. Finally, we investigate the persistent behavior characteristic of the Hurst phenomenon in the distribution of mutations along SINV genomes, finding evidence that, in the sudden treatment, mutations become more randomly distributed earlier in time.

## 2. Methods

### 2.1. Description of the study system and data acquisition

We analyzed data from an experimental evolution study that tracked the molecular adaptation of SINV (species *Alphavirus sindbis*, genus *Alphavirus*, family *Togaviridae*) typically cultured on a highly susceptible host, BHK-21 cells, while challenged to infect the less-susceptible host CHO cells, which were genetically modified (pgsD-677 ATCC CRL-2244) to be more resistant to SINV infection [1]. Briefly, SINV populations were evolved through 25 passages (∼100 virus generations) in monolayer cell cultures with a multiplicity of infection (MOI) of ∼0.01 plaque-forming units (pfu) per cell at each passage. Two different treatments from the original study [1,2] were chosen for an in-depth analysis to investigate the link between population parameters and system dynamics of molecular adaptation to a novel host. The initial stock was prepared by expression of an infectious clone in BHK-21 cells for 24 h [16]. In the gradual treatment, the proportion of CHO cells in the cell cultures increased at each passage, reaching 100% at the last passage. In contrast, in the sudden treatment, cell cultures were entirely composed of CHO cells from the first passage. For each treatment, nine populations were evolved and sequenced at passages 4, 7, 10, 13, 16, 19, 22, and 25. Following RNA extraction, two technical replicates were prepared for each sample from the reverse transcription step onward. Sample preparation is fully explained in [2].

The final allele frequency tables for each sample were obtained from [2]. Briefly, reads were trimmed with cutadapt version 1.8.3 [17], aligned to the consensus sequence of the original stock with BWA version 0.7.10 [18] and variant calling was conducted with QUASR version 7.01 [19] and VarScan version 2.3.9 [20]. Only variants with a frequency higher than 1% and present in both technical replicates were kept.

### 2.2. Wright-Fisher approximate Bayesian computation estimates of selection coefficients and effective population sizes

The selection coefficient (*s*) and effective population size (*N_e_*) were estimated for each allele with approxwf [21], which uses a discrete approximation of the Wright-Fisher diffusion model and Bayesian inference to estimate population parameters. The implementation assumes a log-Uniform(1, 5) prior distribution on *N_e_* and a Normal(0, 0.05) prior distribution on *s* [21]. The algorithm runs an MCMC using 51 states for 25,000 interactions, discarding the first 2000 interactions as burn-in. The mean values of *s* and *N_e_* from their posterior distributions were used in downstream analyses.

### 2.3. Genetic diversity within evolving viral populations

To measure the genetic distance between two viral populations from consecutive time points, the allele frequency difference (AFD) [22] was calculated at each polymorphic site and averaged by the richness of the sample with custom R scripts, where *n* represents the total number of different alleles observed at the polymorphism and the *fi* terms are the proportion of allele *i* in the two viral populations.

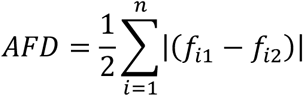

As a second measure of within-sample genetic diversity, we used allele richness, defined as the number of unique alleles per locus, adjusted for sample size.

### 2.4. Taylor’s fluctuations power law

Mean and variance of allele frequencies (*p*) across time (〈*p*〉 and 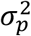, respectively) were computed and used to estimate Taylor’s parameters *V* and *β* by linear regressions in the log-log space: 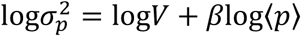 [3]. Given that the proportion of zeros had a significant influence on the fits, Taylor’s parameters were estimated independently for alleles grouped by their number of non-zero occurrences. Multivariate analysis of variance (MANOVA) of Taylor’s parameters were performed with parameters estimated from at least five observations. The magnitude of effects was evaluated using the 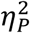 statistic. Conventionally, 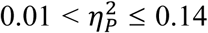 are considered as medium effects and 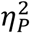 > 0.14 as large ones. Three random walk matrices were constructed based on the cumulative sum of random variables drawn from a Normal distribution with a mean of zero and standard deviations of 0.2, 0.05 and 0.01, respectively. For each matrix, 1000 random walks with eight time points were generated, and values below 0.01 were set to zero and those above 0.99 were set to one. After eliminating walks with only zeros, 792, 757 and 587 random walks were present in the matrices with standard deviation 0.2, 0.05 and 0.01, respectively.

### 2.5. Long-range dependence of mutation sites

To characterize the long-range dependence behavior of mutations along the SINV genome, rescaled-range analyses were performed on the placement of mutations on each time point with the R package pracma version 2.4.2 [23]. For this analysis, binary sequences of length 11,703 (which is the genome length) with ones in the position of mutations were used to estimate the empirical Hurst’s exponent (*H*). The minimum window size was set to 2000 to avoid windows with only zeros. Indels were excluded from this analysis.

### 2.6. Statistical analyses

All the statistical analyses indicated above were performed with R version 4.4.0 under RStudio version 2024.04.2+764.

## 3. Results and Discussion

### 3.1. A sudden host transition is associated to stronger genetic drift

The selection coefficient (*s*) and effective population size (*N_e_*) were estimated for each allele of the evolving viral populations (figure 1). Regardless of the cistron, the sudden host transition treatment consistently resulted in ∼2 times smaller *N_e_* values. Because *s* and *N_e_* can influence each other, we tested for the effects of cistron, treatment, population and their interactions on both *s* and *N_e_* parameters. All independent variables and their interactions, with the exception of the interaction between treatment and cistron, were significant (MANOVA; *P* < 0.0001 for treatment, cistron and population, and *P* = 0.0012 for the interaction between cistron and population). By analyzing *s* and *N_e_* independently, treatment had the relatively greater effect on *N_e_* but very small effect on *s* (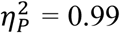 for *N_e_* and 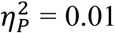 for *s*). However, *post hoc* pairwise tests show that, on average, alleles observed in the sudden transition at the *nsp1*, *nsp2* and *nsp3* cistrons are significantly less deleterious that those found under the gradual transition. The proteins encoded by these three cistrons are involved in the formation of the viral replication complex [24], thus being expected to be under strong purifying selection. In particular, the product of *nsp3* is involved in host specificity and virulence.

**Figure 1.**
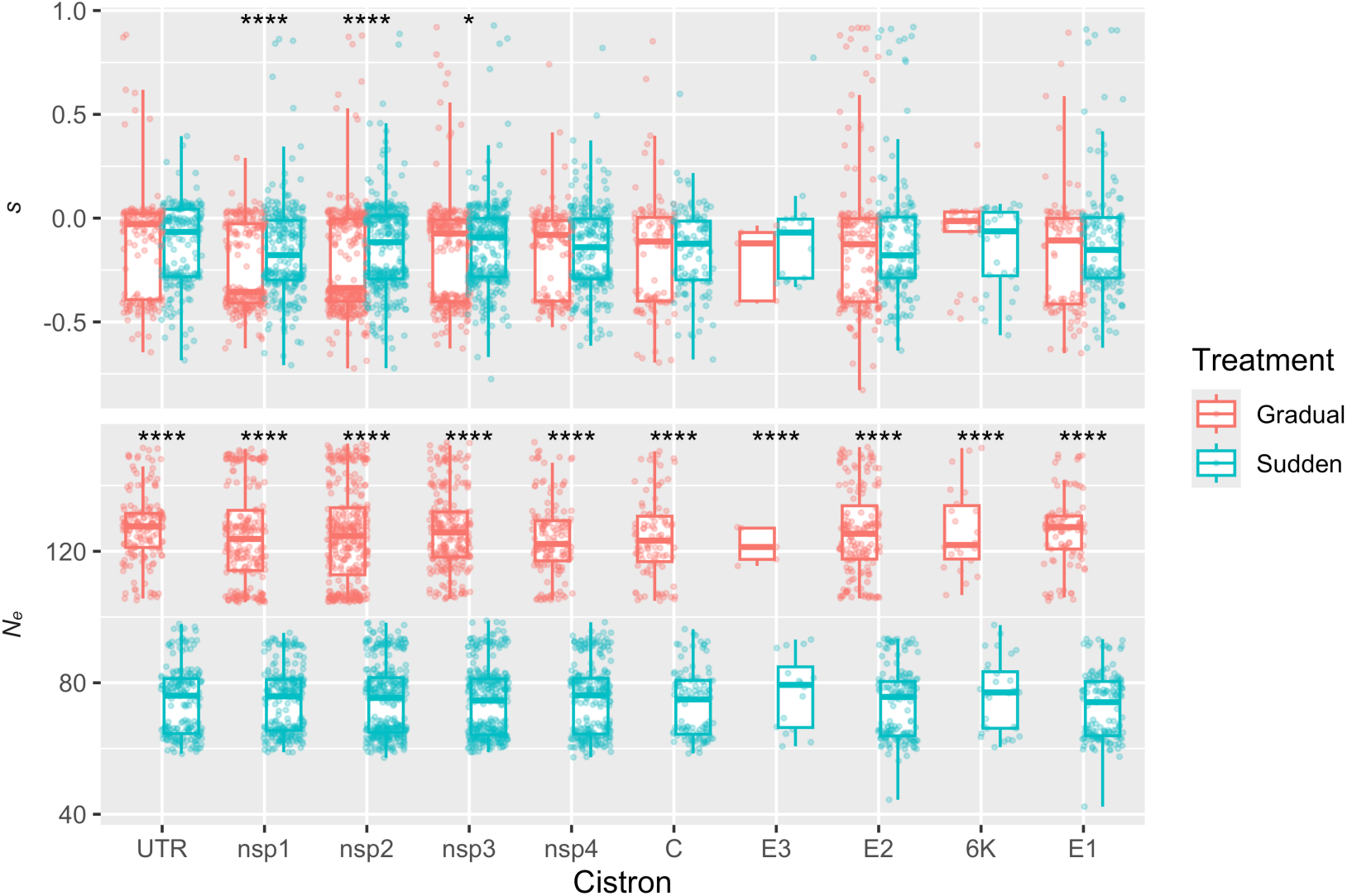
Population parameters *s* and *N_e_* for each allele across all populations. Boxplots of the mean values of the posterior distribution of *s* and *N_e_* for each allele, displayed by treatment and cistron. Asterisks represent the significance of Mann-Whitney two-samples tests: * *P* < 0.05 and **** *P* < 0.0001.

Sudden host transition was associated with stronger (more extreme) bottlenecks, that led to reduced *N_e_*, which in turn increased the effects of genetic drift. Interestingly, in contrast to the gradual treatment, cistron had no effect on *s* (ANOVA; *P* < 0.0001 for the gradual and *P* = 0.3400 for the sudden treatments, respectively). An explanation could be that under the bottleneck the effect of selection across different regions may be weaker or less discernible, while the effect was more apparent when the population was less affected by drift, as in the gradual treatment. However, despite this difference, the effect size of cistron on *s* in the gradual treatment was still moderate 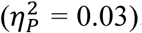.

### 3.2. A sudden host transition is associated to more instability population allele composition

Changes in the composition of the populations between consecutive passages were evaluated by summing the allele frequency differences (AFD) [22] of all alleles. Overall, the composition of the populations was more variable (less steady) in the sudden treatment (figure 2), most likely due to greater effects of genetic drift. In the gradual treatment, most populations changed steadily until the last passage. At this point, a large shift in the composition of the populations was seen (figure 2a).

**Figure 2.**
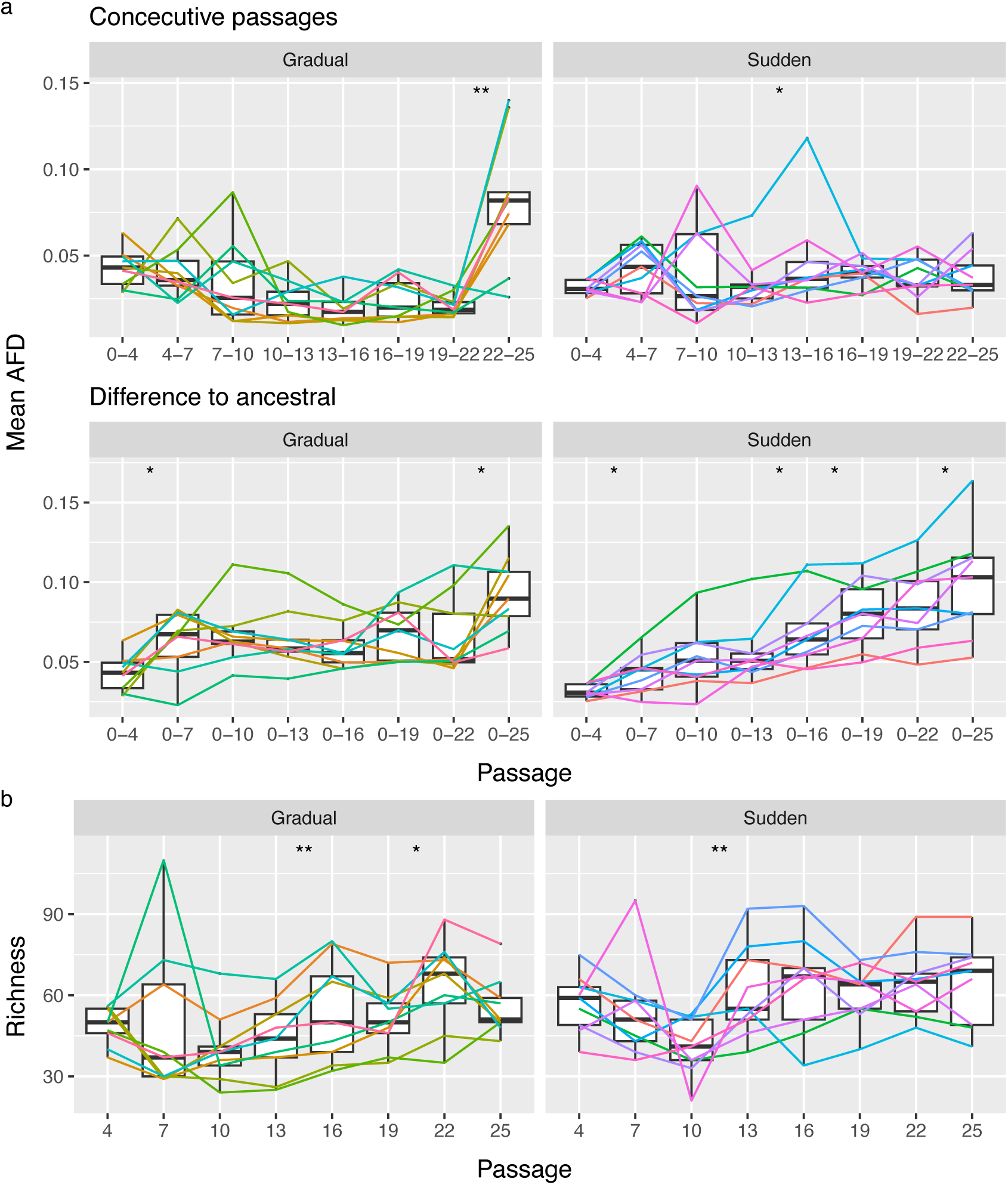
Evolution of the populations’ composition. (a) Boxplots of the mean AFD between two consecutive passages and between each passage and ancestral sequence, also showing the mean AFD for each population represented by lines. (b) Boxplot of the richness of each passage. Lines represent the richness of each population, and asterisks represent the significance of paired samples Wilcoxon tests: * *P* < 0.05 and ** *P* < 0.01.

The rate at which the populations diverged from the ancestral sequence also differed between treatments (figure 2a). While in sudden treatment a trend of constant differentiation from the ancestral is seen, in the gradual treatment, we have that, at some points, populations do not diverge from, or even become more similar to, the ancestral. This phenomenon, known as leapfrog effect [25], happens when one haplotype supersedes another and both have evolved from the same common ancestor. Since both haplotypes are more similar to their ancestor than they are to one another, the population can become more similar to the ancestral population as it evolves. Five not mutually exclusive mechanisms can be brought forward to explain this leapfrog effect. First, balancing selection that might favor diversity at certain loci, allowing rare haplotypes to gain an advantage under changing environmental or selective conditions [26]. Second, epistasis and hitchhiking in which specific combinations of alleles in rare haplotypes may confer a fitness advantage (epistasis) that becomes beneficial over time, hitchhiking them to dominance [27–29]. Third, environmental fluctuations can alter the fitness landscape, favoring haplotypes that were previously at a disadvantage, even if they are less related to the dominant haplotypes [30–32]. Fourth, genetic drift in small population allows rare haplotypes to randomly increase in frequency, contributing to their rise in dominance over time [33]. And, fifth, lineage sorting, by which low-frequency alleles unrelated to previously dominant ones but linked through deeper evolutionary events [34], and recombination that breaks apart haplotypes and creates new advantageous combinations that are not present in the previously dominant group [35]. Since we have observed the leapfrog phenomena in the gradual transition regime (larger *N_e_*; figure 1), we propose that genetic drift may contribute less than the other four mechanisms.

### 3.3. BHK-21-specialists persisted until the complete elimination of BHK-21 cells from the environment

We hypothesize the co-occurrence of BHK-21-specialist, CHO-specialist and generalist haplotypes in the gradual treatment. Clonal interference and competition are expected between and within generalists and specialists, but less so between haplotypes specialized in different host cells. Clonal interference occurs when beneficial mutations are lost due to the competition with other beneficial mutations that are present in other haplotypes [25,36,37]. Here, it is more likely that beneficial mutations on generalists will be lost due to competition with specialists, although the opposite may also happen, especially if there is no tradeoff in the evolution of generalists [38]. The large shift in population composition seen at the last passage in the gradual treatment suggests that BHK-21-specialists became extinct once BHK-21 cells were completely absent from the environment. We noted, however, that two populations did not seem to follow this trend, at least as strongly, suggesting that in those cases, a large fraction of high-fitness generalists could have emerged (figure 2a).

To better understand the evolution in the composition of the populations, the richness of each sample, defined as the number of polymorphic sites, was analyzed (figure 2b). In particular, we were interested to investigate whether the elimination of BHK-21-specialists was, as expected, associated with a loss in richness. Interestingly, we noted that in the sudden treatment, a loss of richness is initially seen followed by a relatively high richness that is maintained from passage 13 onward. This observation, together with the fact that at the same time populations are evolving with high variability, suggests that, between passages, there is a high number of mutations that are lost but are being replenished by new ones. Focusing again on the gradual treatment, while a drop in richness can be seen at the last point, it is not significant. Thus, we found that the apparent loss of BHK-21-specialists does not fully explain the large shift in population composition observed at the last passage in the gradual treatment.

We reanalyzed data of the fitness of the evolved populations on BHK-21 and CHO cells to better contextualize the results obtained above (figure 3). The fitness of populations evolved in the sudden treatment is higher in CHO cells than in BHK-21 cells, consistent with populations majorly composed by CHO-specialist haplotypes. However, no significant difference in fitness was seen for populations evolved in the gradual replacement regime. Given that there is no tradeoff between hosts in these populations [1], our results suggested that, for the gradual treatment, the fitness of CHO-specialists plus generalists in CHO cells is similar to the fitness of generalists in BHK-21 cells. Thus, gradual host replacement is associated not only with the emergence of generalist populations but also of high-fitness generalist haplotypes [1].

**Figure 3.**
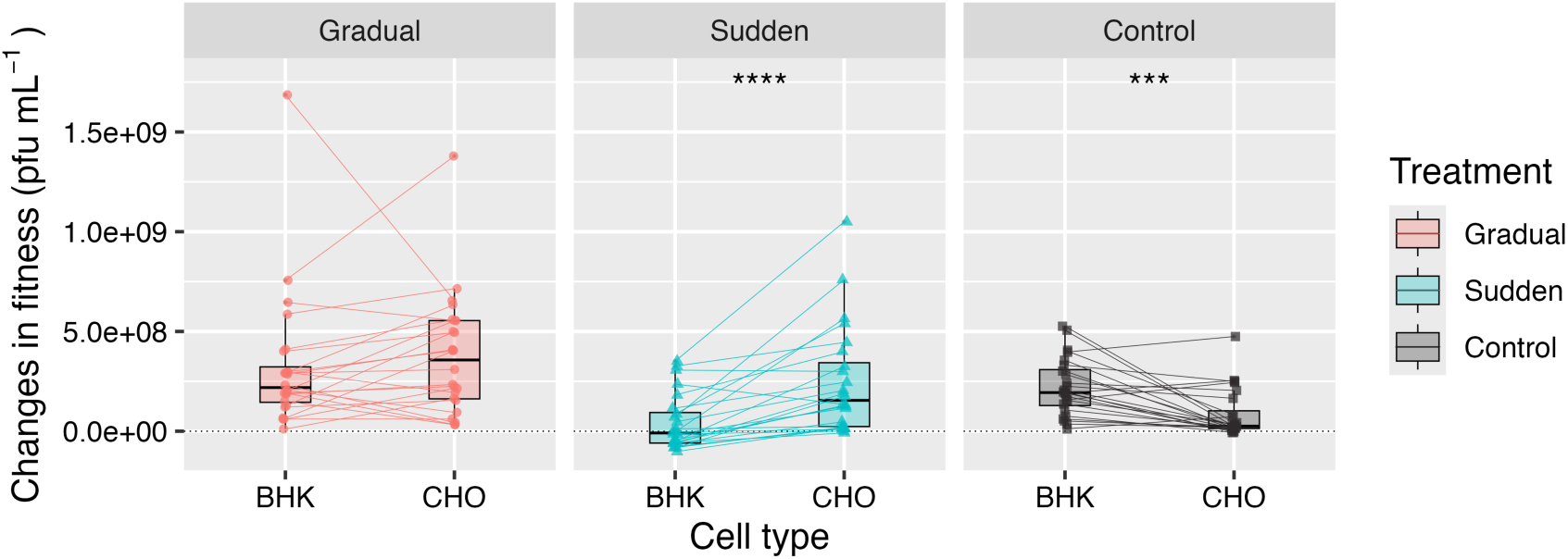
Effect of host replacement regime on fitness. Boxplots of the change in fitness relative to the ancestral for control (lineages evolved in BHK-21 cells), sudden and gradual treatments. Lines represent lineages, and asterisks represent the significance of paired Wilcoxon tests: *** *P* < 0.001 and **** *P* < 0.0001.

### 3.4. Allele frequencies display large fluctuations and aggregation behavior

Morley et al. [2] observed that some positively selected mutations were lost after reaching intermediate allele frequencies, suggesting a clonal interference effect, which was particularly pronounced in the gradual treatment lineages. Here, we sought to determine whether temporal fluctuations in allele frequencies followed Taylor’s power law. We hypothesize that due to the larger effect of drift on smaller populations, alleles in the sudden treatment will fluctuate closer to what is expected by the Poisson distribution, and thus, present smaller *β* than alleles in the gradual treatment. Additionally, large fluctuations due to clonal interference, especially for those alleles that reached intermediate or high frequencies, are also expected in the gradual treatment. Inspection of the Taylor’s plots showed that the fraction of zeros had a drastic impact on the fit (figure 4), which is to be expected given that only eight time points are present. Here, it is important to note that the presence of zeros might be either due to biological reasons, *i.e*., the allele was not present at that time point, or due to technical noise (measurement error). The latter is more likely for low frequency alleles that fluctuate close to the detection limit.

**Figure 4.**
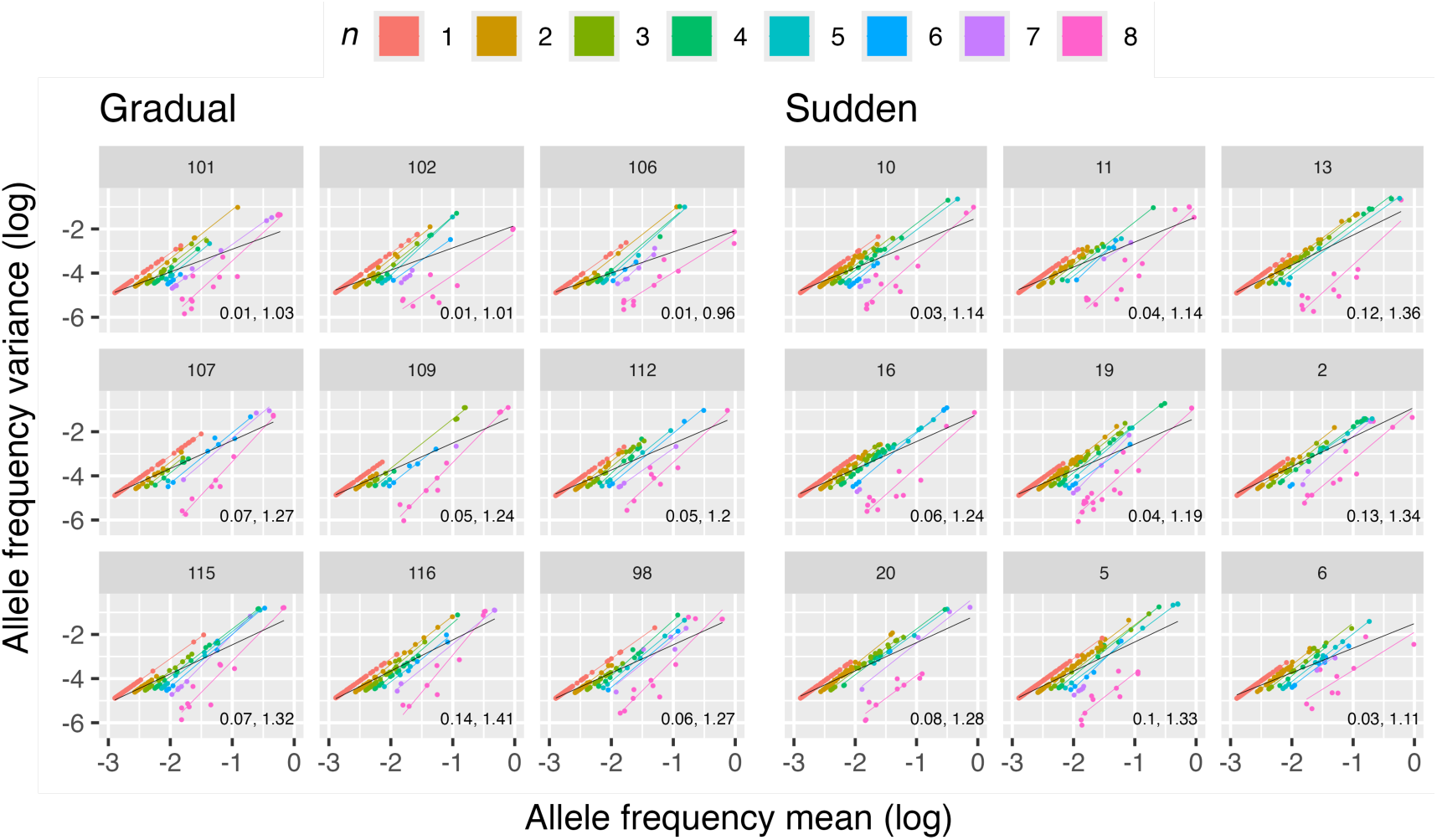
Taylor’s law plots. Each panel corresponds to one population, where each point corresponds to the mean and standard deviation of an allele frequency across all passages. Alleles are colored by their number of non-zero occurrences (*n*), and for each *n*, a colored line represents the fit to Taylor’s law. A black line represents the fit to Taylor’s law for all alleles together. Taylor’s parameters *V* and *β*, respectively, for the fit without adjusting for *n*, are shown.

Fits to Taylor’s law were then performed using alleles grouped by their number of non-zero occurrences (*n*) to analyze Taylor’s parameters *V* and *β*, henceforth *V_n_* and *β_n_*, and how they scale by host-change regime. In all cases, *β_n_* was estimated to be close to or higher than one (figure 5a). When performing fits by the number of zeros, we expect that, as the mean allele frequency increases, we tend to observe aggregation since values will be distributed across non-zero spaces. However, parameter *β_n_* is still able to capture how these values are distributed and measure the strength of aggregation, with an important caveat. Due to the bounded measures of allele frequency in the interval (0, 1), early fixation of an allele will make it so that it will exhibit high mean frequency and relatively low standard deviation, given that most time points will be ones. This, in turn, will drive *β_n_* down, making it to resemble a Poisson distribution. Late fixation will conversely drive *β_n_* up. Thus, this parameter must be analyzed in respect to its deviation from 2, which is the expected for random fluctuations without fixation. In other words, mean frequencies and their fluctuations will occupy different regions in respect to *β_n_* = 2 (figure S1a)

**Figure 5.**
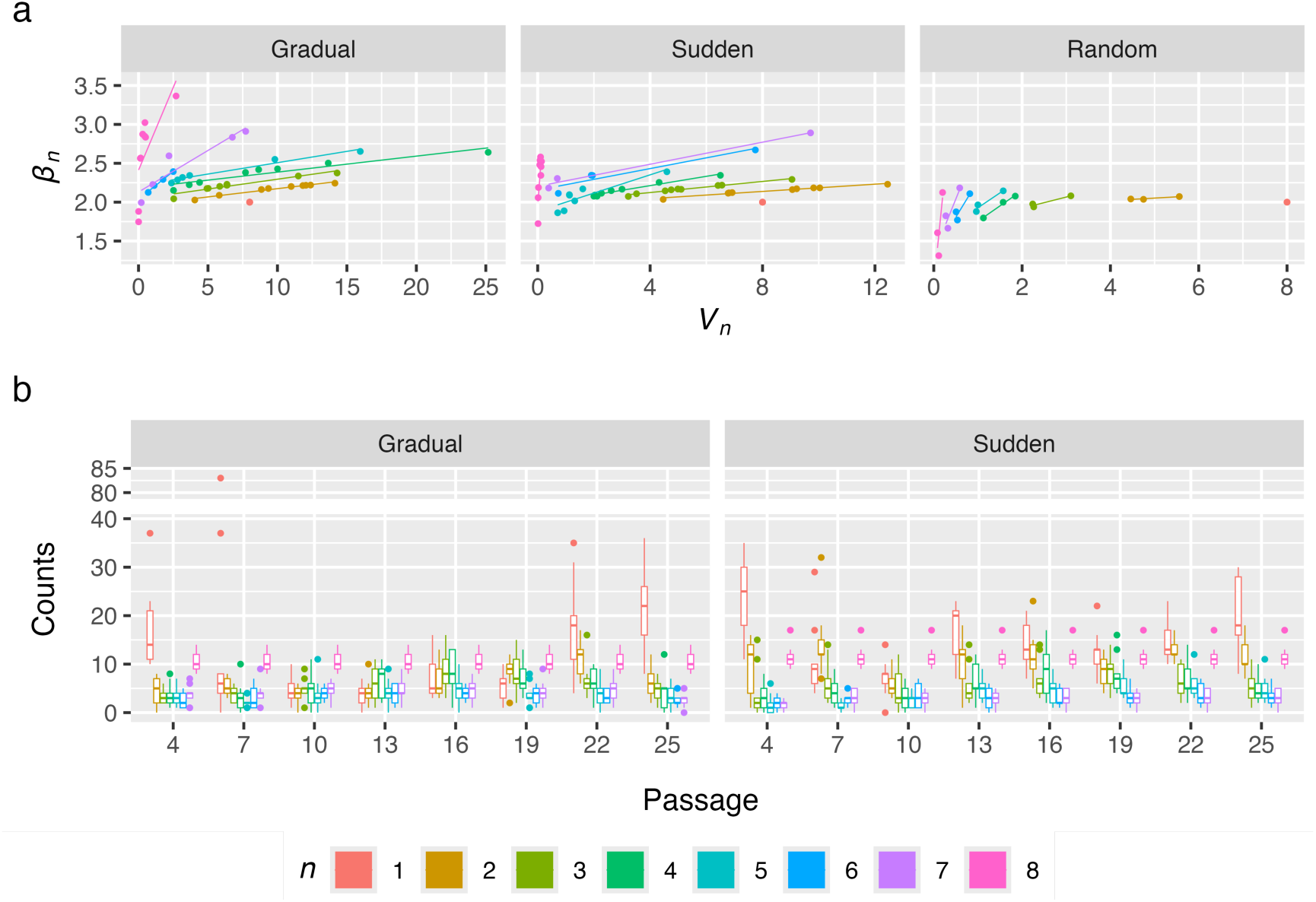
Distribution of alleles frequencies across passages by their *n*. (a) Taylor’s parameters estimated from alleles frequencies for each *n* for the gradual and sudden treatments and random walk matrices. (b) For each passage, boxplots of the distribution of alleles at each category of *n*.

To test this, three simulated matrices of random walks were generated based on the cumulative sum of random variables drawn from a Normal distribution with mean zero and standard deviations of 0.2, 0.05 and 0.01, respectively (figure S1b), where the higher the standard deviation, the higher the fluctuation between time points. We then performed fits to the simulated random walks and observed that, in this case, *β_n_* tends to be lower than one when the frequency of zeros reduces (figure 5a). Additionally, *β_n_* decrease as fluctuation increases, indicating that higher fluctuations are associated to lower *β_n_*. This is due to alleles occupying different regions of the Taylor’s plot depending on the strength of their fluctuations (figure S1c). As expected, when fitting to Taylor’s law without adjusting for *n*, random matrices behave close to a Poisson distribution (*β* ∼ 1), where parameter *V* increases with larger fluctuations.

To better understand whether frequencies are distributed across the sequenced passages, the number of alleles separated by *n* at each passage and plotted (figure 5b). In the sudden treatment, a higher concentration of alleles with an *n* of one or two at the first passages is detected, in agreement with a strong bottleneck at the initial passages as the virus populations are introduced to a completely new host CHO. The number of alleles with an *n* of one or two starts to rise again in the sudden treatment at passage 13; and overall, a higher number of alleles with an *n* of one or two across all passages is seen in comparison with the gradual treatment (figure 5b). This is in accordance with the expected effects of genetic drift, and also with the continuous emergence and loss of new mutations in these shifting populations that was discussed above. In the gradual treatment, alleles with an *n* of one or two are also concentrated at the first and last passages. However, in comparison to the sudden treatment, they are less concentrated at the first passages and their number only starts to rise again at passage 22 (figure 5b).

### 3.5. Effects of treatment and selection coefficient on alleles’ fluctuation

Next, we tested the effect of treatment, population and *n* on the distribution of alleles accounting for all interactions between these variables. The number of non-zero occurrences *n* had the highest influence on Taylor’s parameters *V_n_* and *β_n_* (MANOVA; *P* < 0.0001; 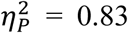). Both the main effect treatment and the interaction between treatment and *n* had a significant influence on Taylor’s parameter (MANOVA; respectively, *P* = 0.0019 and 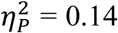, and *P* = 0.0080 and 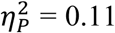), which indicates that host-change rate has a significant and large impact on the distribution of allele frequencies. The main effect population and the interaction between *n* and population were not significant, despite large effect sizes (MANOVA; respectively, *P* = 0.0531 and 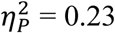, and *P* = 0.0837 and 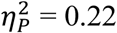).

Alleles were divided into three categories, neutral, deleterious and beneficial, to determine whether strength of selection has an impact on Taylor’s parameters. By visualizing the distribution of *s* (figure 6a) it becomes clearer that *s* follows a mostly bimodal distribution with a peak for negatively selected alleles and another for neutrally selected ones around zero, with only a fraction being positively selected. Interestingly, negatively selected alleles on the gradual treatment seem to be subjected to stronger negative selection. Based on these distributions, we set a threshold of *s* < −0.16 for alleles to be considered as negatively selected and *s* > 0.16 for them to be considered positively selected, whereas alleles with *s* ϵ (−0.16, 0.16) are considered as neutral.

**Figure 6.**
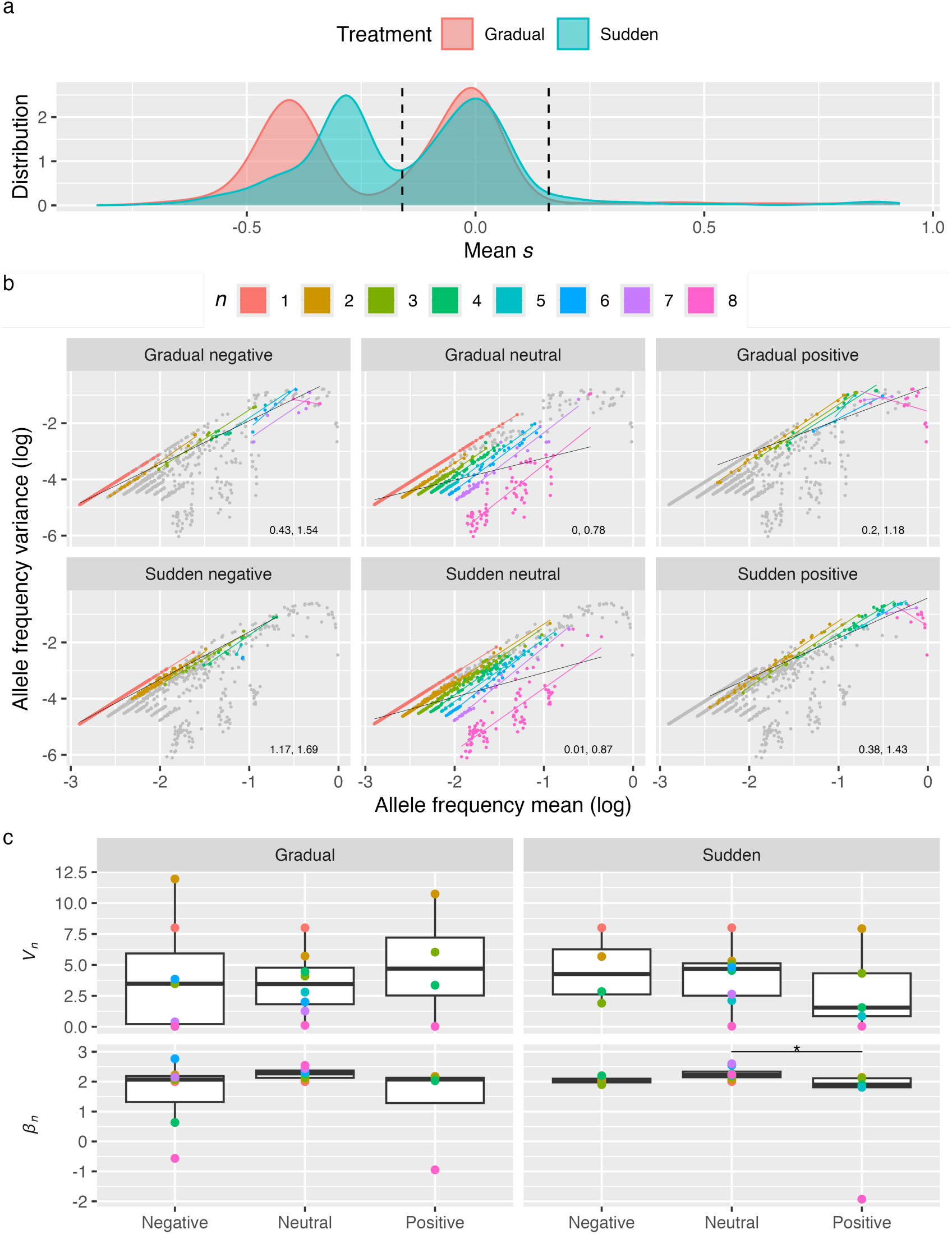
Effects of selection on allele fluctuations. (a) Distribution of *s* per treatment. Dashed lines represent the thresholds for categorizing neutral, negatively and positively selected alleles. Taylor’s parameters *V* and *β*, respectively, for the fit without adjusting for *n*, are shown. (b) Taylor’s plots of combined allele frequencies across populations by treatment and *s*. All the alleles of each treatment are represented as dots and are colored by their number of non-zero occurrences (*n*) in their selection classification. For each *n* and *s*, a colored line represents the fit to Taylor’s law. A black line represents the fit to Taylor’s law for all alleles classified as the corresponding *s* in the panel. Taylor’s parameters *V* and *β*, respectively, for the fit without adjusting for *n*, are shown. (c) Boxplots of *V_n_* and *β_n_* by treatment and *s* category. Asterisks represent the significance of unpaired Wilcoxon tests: * *P* < 0.05.

Given that the separation of alleles by *s* drastically reduces sample size in some populations and *n* combinations, fits to Taylor’s law were performed by treatment, (where frequencies from all population were combined), selection and *n* (figure 6b). Still, sample size drastically affected the fits, especially in cases where alleles with low mean were not present (*e.g*., positively selected alleles with *n* = 8) to accurately scale how standard deviation varies with the mean. Additionally, some *n* categories are missing entirely for either positively or negatively selected alleles depending on treatment. Again, *n* had the highest influence on Taylor’s parameters *V_n_* and *β_n_* (MANOVA; *P* < 0.0001; 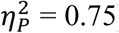), followed by the interaction between *n* and *s* (*P* = 0.0011; 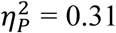), and *s* (*P* = 0.0299; 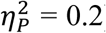). Here, treatment did not have a significant impact on Taylor’s parameters. It is likely that by combining the allele frequencies across populations the analysis became confounded. This is supported by the fact that the interaction between *n* and population was significant when Taylor’s parameters were estimated separately for each population.

Despite not finding a significant influence of treatment on Taylor’s, some very interesting observations can be made from these Taylor’s plots in regard to *s*. Firstly, points are far more dispersed for neutral alleles, and occupy a region composed by alleles that are present at several time points but have low mean frequency and low standard deviation. Most notably, all alleles with *n* = 8 that have low mean frequency and standard deviation are neutral. Due to this, if the data is fitted to Taylor’s law without adjusting for *n*, neutral alleles will be the ones closer to a Poisson distribution. Also, neutral alleles exhibit the highest values of *β_n_* (figure 6c). Secondly, positively selected alleles with an *n* = 8 have negative *β_n_* and their standard deviation decreases as their mean frequency increase. This is due to early fixation, causing most time points to be ones, and consequently, their standard deviation will be low. When fitting the data to Taylor’s law without adjusting for *n*, this drives *β* down, and as a result, *β* is lower for positively selected alleles in comparison with negatively selected ones. Additionally, positively selected alleles occupy a region of the Taylor’s plot close to the upper limits of the system, especially for higher *n*. Thirdly, negatively selected alleles occupy a region of the Taylor’s plot between neutral and positively selected alleles. This difference is clearer for the sudden treatment.

### 3.6. Host replacement regime affects the distribution of mutations along SINV genome

Lastly, it is likely that host change dynamics may also have an effect not only on the temporal aggregation dynamics of allele frequencies but also on genomic location of mutations due to the interplay between drift and selection and linkage. Due to genetic hitchhiking [2,27–29], neutral or low fitness mutations may benefit from being strongly linked to high fitness ones. In addition, the gradual host replacement is associated with greater clonal interference [2], in which genomes containing multiple beneficial mutations outcompete those containing only one, promoting the fixation of linked mutations [2,39–41]. Given that the fixation of sweeps of mutations is greater in the gradual treatment [2], we sought to investigate whether mutations in the gradual treatment are less evenly distributed along the genome, meaning if they are aggregating.

For this, we estimated the empirical Hurst’s exponent *H* to measure persistent behavior in the placement of mutations at each passage for each population (figure 7). Here, *H* was estimated from binary sequences with the same length as SINV genome, where ones represent sites with any disagreement to the ancestral sequence. Despite high variability in the estimation, especially for the sudden treatment, evidence of persistent behavior was found in most cases, were the median value at each passage for both treatments were always above 0.5 (figure 7). At the first two initial passages, median *H* is higher for the sudden treatment, follow by two passages where *H* is more or less similar between treatments, and finally, *H* decreases and becomes smaller on the sudden treatment (despite these differences are only significant in passage 22, *P* = 0.040). This suggests that the continuous emergence and loss of mutations on the sudden treatment that was previously observed at the final passages is associated with more random scattering of mutations along the genome, and might be a big factor on why mutations are more randomly positioned on this treatment. However, we found that treatment did not have a significant influence on *H* (linear model with passage and treatment as orthogonal fixed effects and population as a random effect nested within treatment; sudden coefficient estimate = –0.043, *P* = 0.100).

**Figure 7.**
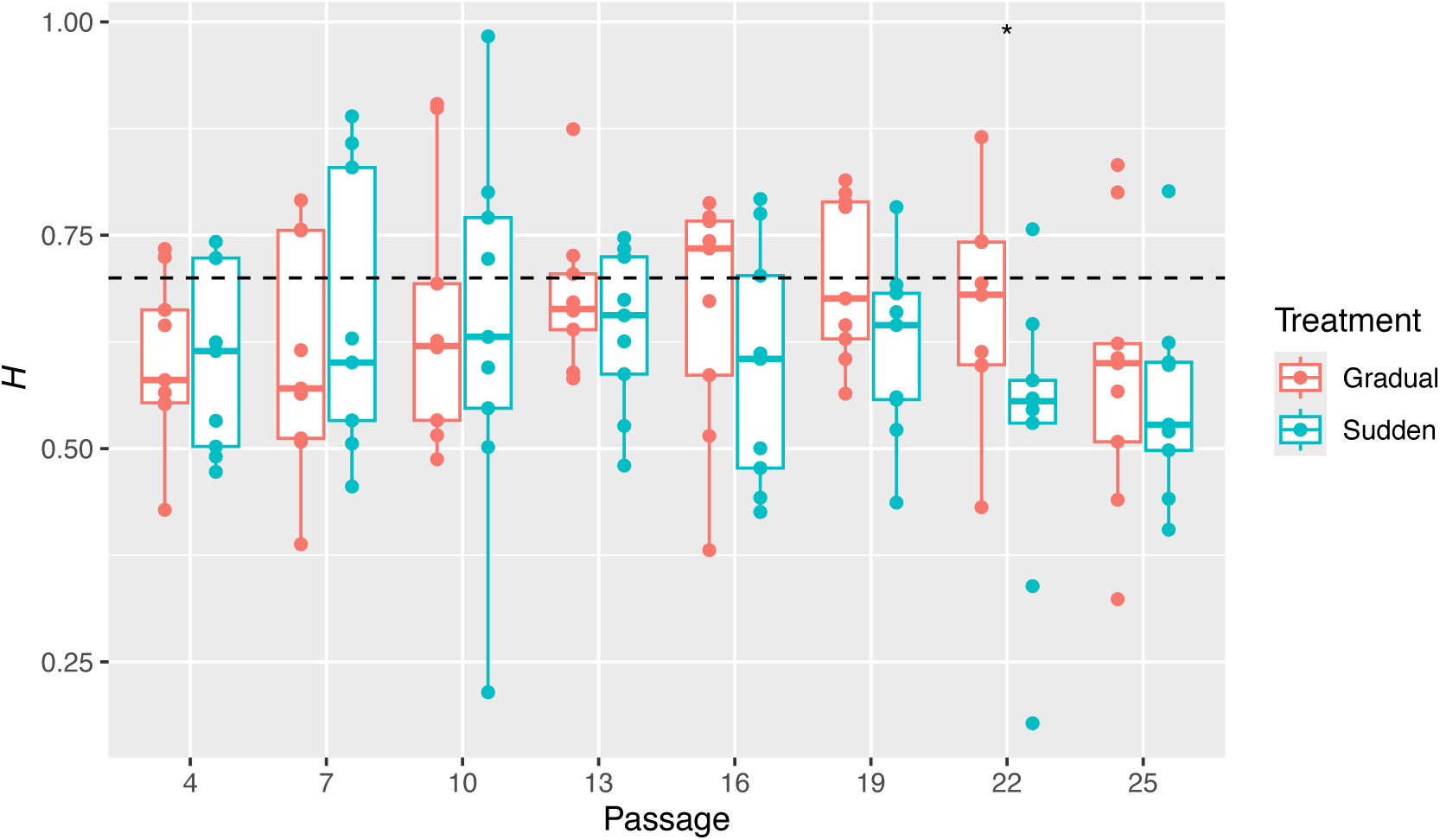
Persistent behavior in the placement of mutations. Hurst exponents (*H*) estimated from binary sequences (where 1 represent a mutant allele) for each population at each passage. Dashed line represents the 0.7 Hurst phenomenon threshold. Asterisks represent the significance of unpaired Wilcoxon tests: * *P* < 0.05.

## 4. Conclusions

This study explores the molecular dynamics of virus adaptation to novel hosts. By reanalyzing the genomic data from experimental evolution of SINV in populations subjected to sudden versus gradual host replacements, we investigated how environmental heterogeneity, genetic drift, and selection influenced temporal fluctuations in allele frequency and genome-wide mutation distribution. Using two mathematical tools from the complex systems toolbox, Taylor’s power law and the Hurst exponent, we characterized whether the observed fluctuations were consistent with random noise or governed by more complex underlying laws.

Our findings emphasize the critical role of environmental heterogeneity in shaping viral adaptation. Gradual host replacement enables more stable and convergent evolutionary trajectories, enhancing viral fitness across environments. This regime promotes clonal interference, facilitating the fixation of linked beneficial mutations and time-persistent genomic mutation patterns. Conversely, sudden host transitions reduce effective population sizes, amplifying genetic drift, destabilizing allele frequencies, and causing more scattered mutation distributions along the viral genomes. These results underscore the importance of the rate and nature of environmental change in determining evolutionary outcomes.

Additionally, applying Taylor’s power law and the Hurst exponent provided insights into the scaling behavior and temporal aggregation of allele frequency fluctuations. Neutral alleles exhibited greater variability, while positively selected alleles tended to fix earlier. The interaction of genetic drift, selection, and linkage influenced mutation distribution, with gradual host replacement favoring aggregated mutation patterns due to hitchhiking of linked mutations.

This study advances our understanding of virus evolution and provides a novel framework for predicting pathogen adaptation to changing environments, such as shifts in host availability or immune pressures.

## Ethics

This work did not require ethical approval from a human subject or animal welfare committee.

## Data accessibility

Supplementary figure S1 (pdf) and the results of approxwf and alleles frequencies (Excel file) are available at the Zenodo repository https://doi.org/10.5281/zenodo.14557803. R scripts used to perform the analyses are available at https://www.github.com/jmfagundes/SINVevo.

## Declaration of AI use

We have not used AI-assisted technologies in creating this article.

## Authors’ contributions

J.M.F.S.: conceptualization, formal analyses, investigation, methodology, software, visualization, writing – original draft, writing – review and editing; M.J.O-U.: conceptualization, investigation, methodology, writing – review and editing; V.J.M.: data curation, investigation; P.E.T.: data curation, investigation, supervision, writing – review and editing; S.F.E.: conceptualization, formal analyses, investigation, methodology, writing – original draft, writing – review and editing.

All authors gave final approval for publication and agreed to be held accountable for the work performed therein.

## Conflict of interest declaration

We declare we have no competing interests.

## Funding

This work was supported by grant PID2022-136912NB-I00 funded by MCIN/AEI/10.13039/501100011033 and by “ERDF a way of making Europe” and by Generalitat Valenciana grant CIPROM/2022/59 to S.F.E. M.J.O.U. was supported by grant FPU2019/05246 funded by MCIN/AEI/10.13039/501100011033 and by “ESF investing in your future”.

## Acknowledgements

The authors would like to thank Dr. Carlos P. Garay and Dr. José A. Oteo for very fruitful discussions on the meaning of Taylor’s power law and Hurst’s exponent in evolving complex systems.

## Notes

https://doi.org/10.5281/zenodo.14557803

https://www.github.com/jmfagundes/SINVevo

### Competing Interest Statement

The authors have declared no competing interest.

### Summary of Updates

Text has been revised in response to the comments received from three reviewers. Text has been expanded in several sections to provide better explanations. Figures have been updated.

