## Supplementary Figure S1 for "Taylor’s power law rules the dynamics of allele frequencies during viral evolution in response to host changes" (from another version of this preprint: version 1, 2024-12-27)

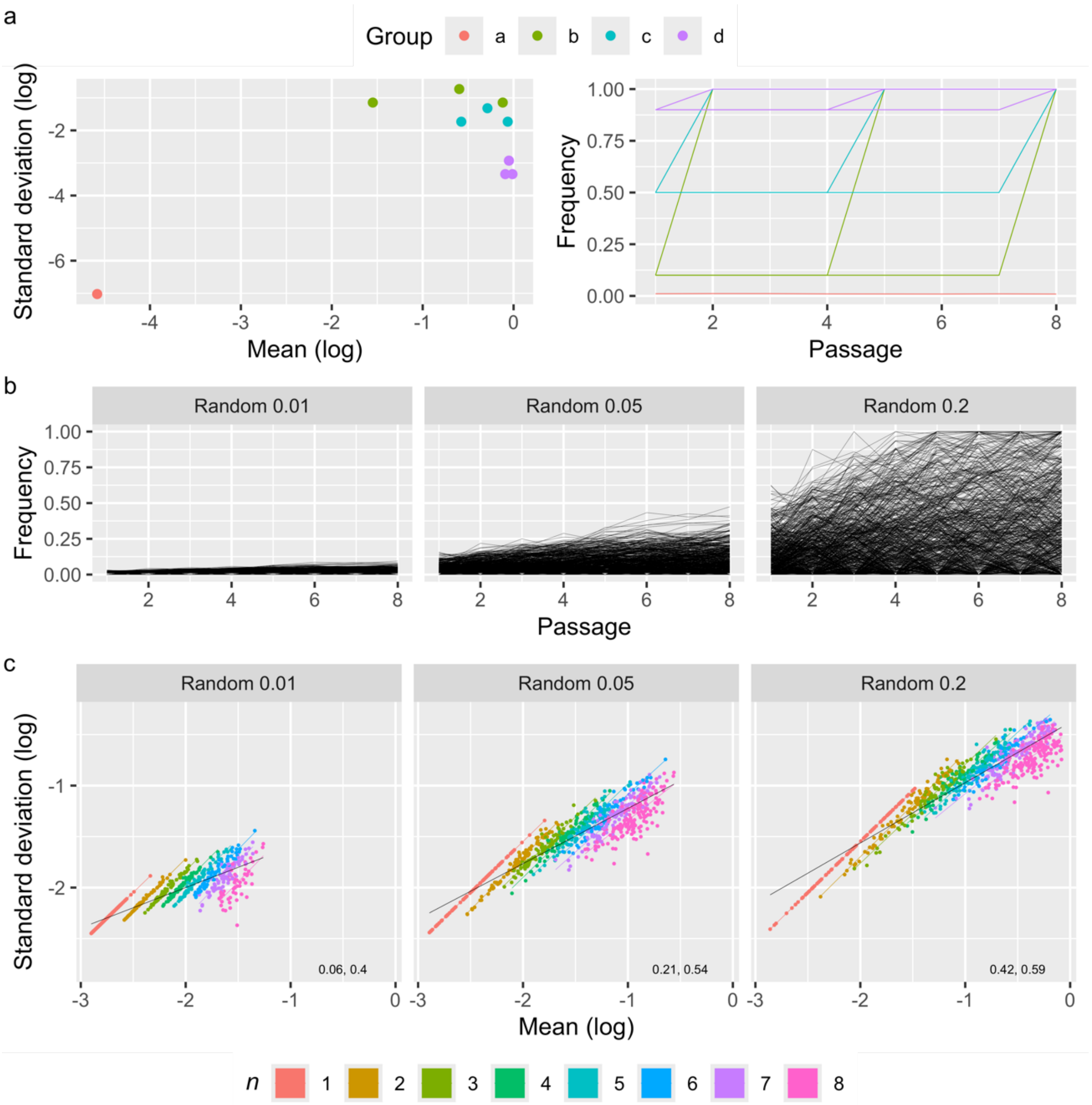


**Supplementary Figure S1.** (a) Mean and standard deviation and trajectories of simulated allele frequencies, where $n = 8$ for all cases, showing how $\beta_n$ scales from alleles with small fluctuations around 0.01 (group a) to large fluctuations (group b), medium fluctuations (group c) and early rise with small fluctuations from 1. (b) Trajectories of random walks from the random matrices built with standard deviation of 0.01, 0.05 and 0.2. (c) Taylor's plot of random matrices, colored by $n$. For each $n$, a colored line represents the fit to Taylor's law. A black line represents the fit to Taylor's law for all alleles. Taylor's parameters $V$ and $\beta$, respectively, for the fit without adjusting for $n$, are shown.
